# The within-person association of relative left frontal activity and vagally mediated heart rate variability not moderated by history of depression

**DOI:** 10.1101/2023.07.10.547869

**Authors:** Yaohui Ding, John J.B. Allen

## Abstract

Motivated by the Neurovisceral Integration Model (NVI) of cardiac vagal control, we investigated the relationship between relative left frontal activity (rLFA) and vagally mediated heart rate variability or respiratory sinus arrhythmia (RSA) in 287 participants, half of whom had a history of depression. We hypothesized that there would be a within-person association of rLFA and RSA such that when RSA is lower rLFA would also be lower (Hypothesis I). Moreover, it was hypothesized that this within-subject association would be moderated by a history of depression (Hypothesis II). Metrics of rLFA and RSA were derived from concurrent electroencephalogram and electrocardiogram recordings. The logarithmic difference in EEG alpha power between the homologous right and left electrodes (Ln (Right/Left)) in the frontal region was used to index rLFA. A Hilbert transform was applied to the mean-centered and bandpass-filtered (0.12-.40 Hz) inter-beat interval (IBI) time series to get a fine-grained measure (in the time domain) of RSA. A linear *mixed ANOVA* model with rLFA as the dependent variable and RSA as the main fixed effect found that participants had less rLFA during epochs when they had lower RSA, which was consistent with the prediction from Hypothesis I. Contrary to the prediction from Hypothesis II, the within-person association of RSA and rLFA was *not* moderated by a history of depression. However, the association between RSA and rLFA varied across the four pairs of frontal electrodes that we examined. Thus, more research is needed to determine the spatial extent of this association, e.g., examining the relationship between source-localized rLFA and RSA.

## 1 | INTRODUCTION

Major depressive disorder (MDD) is a debilitating and prevalent neuropsychiatric condition (Kessler and Bromet, 2013) that places a heavy burden on individuals and incurs a great economic cost to society (Greenberg et al., 2015). According to estimates from a Global Burden of Diseases study, MDD was the 4th leading cause of disability in 1990 and will be the 2^nd^ leading cause of disability in 2020 (Ferrari et al., 2013). As a result, MDD is one of the most researched topics in psychiatry, clinical psychology, neuroscience, and psychophysiology, with the focus on finding risk factors for, mechanisms of, and ultimately effective treatments for MDD. With respect to the research on MDD within the field of psychophysiology, relative left frontal activity (rLFA) (Reznik and Allen, 2018; Thibodeau et al., 2006) and vagally mediated heart rate variability (Bassett, 2016; Beauchaine, 2015) have been investigated extensively over the last few decades for their relationships to MDD specifically and psychopathology in general. Both measures have been related to a history of depression or risk for depression (Allen and Reznik, 2015; Kemp et al., 2010; Reznik and Allen, 2018), yet no research has examined how these measures relate to each other within person, and whether this relationship is moderated by a history of depression. In the present study, we investigated the relationship between relative left frontal activity and RSA using concurrent EEG and EKG recordings from 287 subjects, about half of whom had a history of depression.

### 1.1 | Relative Left Frontal Activity and History of Depression

Relative left frontal activity is a metric derived from the logarithmic difference in the power of alpha band EEG activity between the homologous **right** and **left** electrodes (Ln(Right)-Ln(Left)) in the frontal region (Allen and Reznik, 2015). This metric has high internal consistency and good test-retest reliability (Allen et al., 2004; Towers and Allen, 2009). It has been proposed as a neurophysiological marker of depression vulnerability (Allen and Reznik, 2015). Specifically, a lower rLFA score, which putatively reflects relatively less left than right cortical activity (Jensen and Mazaheri, 2010; Klimesch et al., 2007; Mathewson et al., 2011), has been associated with a history of or a risk for depression (Harmon-Jones and Allen, 1997; Stewart et al., 2010; Vuga et al., 2006). On the contrary, a higher rLFA score is associated with psychological well-being and effective emotion regulation (Davidson, 2004; Jackson et al., 2003). The reviewers thought this section was too concise.

### 1.2 | Vagally Mediated Heart Rate Variability and Major Depressive Disorder

Vagally mediated heart rate variability is heart rate variability in the high frequency band (0.12 – 0.4 Hz), which can be used to index parasympathetic efferent activity to the heart (Porges, 2007). To date, numerous studies have investigated its relationships to stress and health (Porges, 1995; Thayer et al., 2012), emotion regulation and social engagement capacity (Porges, 2001, 2007; Porges et al., 1994), depression and several other forms of internalizing psychopathology (Beauchaine, 2015; Chalmers et al., 2014). In depression, a lower RSA at rest has often been observed (Bassett, 2016; Kemp et al., 2010; Koch et al., 2019; Rottenberg, 2007).

There has been a long-standing interest in delineating the anatomical connections and functional integration between the central nervous system and visceral organs, particularly the heart, in health and disease. The dynamic change in heart rate is a result of the complex interactions among intrinsic cardiac mechanisms (Jose and Collison, 1970), sympathetic and parasympathetic innervations of the pacemaker cells on the heart (Levy, 1990). Sympathetic input to the heart acts slowly but increases the heart rate, whereas parasympathetic input acts quickly but decreases the heart rate. Craig (2015) argued that both the efferent innervation of the heart and afferent activity from the heart are asymmetric and that these asymmetries are respected in the central nervous system and proposed a homeostatic neuroanatomical model of forebrain emotional asymmetry. According to this model, parasympathetic cardiac activity is predominantly influenced by the left hemisphere and sympathetic activity by the right hemisphere (Craig, 2005). Thus, hypoactivity of the left frontal cortex, which is often observed in subjects with a history of depression, leads to increased heart rate and decreased vagally-mediated heart rate variability.

### 1.3 | Present Study & Hypotheses

The relationship between relative left frontal activity and RSA within-persons has not been investigated in the context of major depression. We set out to test this relationship empirically using simultaneous EEG and EKG recordings from 287 subjects, about half of whom have had a history of depression. Motivated by the homeostatic neuroanatomical model of forebrain emotional asymmetry, we hypothesized that there would be a within-subject association of relative left frontal activity and RSA such that when RSA is lower, relative left frontal activity would be lower, reflecting relatively less left frontal activity (**Hypothesis I**). Moreover, because both lower rLFA and RSA scores have been observed in depression, we hypothesized that the within-subject association of rLFA and RSA would be weaker in subjects who have had depression (**Hypothesis II**).

## 2 | METHOD

### 2.1 | Participants

The present sample is derived from the sample reported in (Stewart et al., 2010), where 306 participants were recruited from courses at the University of Arizona and the local community in Tucson, Arizona. Beck Depression Inventory (BDI) (Beck et al., 1996) scores were used to identify prospective participants. To obtain a more representative sample, individuals with BDI scores across the entire range of depression severity (range=0-45.5, mean=10.9, SD=10.4) and a broad range of symptoms (no symptoms, few symptoms, several symptoms, and full clinical severity) were invited for screening. Participants were screened for exclusionary criteria during both the phone interview by a post-baccalaureate project manager and the intake interview conducted by graduate- or post-graduate-level clinical raters. Exclusionary criteria included left-handedness, history of head injury with loss of consciousness >10 minutes, concussion, epilepsy, electroshock therapy, use of current psychotropic medications, any current comorbid DSM-IV Axis I disorder other than lifetime MDD or current dysthymia in the absence of a history of MDD, as determined by the Structured Clinical Interview for DSM-IV (American Psychiatric Association 1994). Active suicidal ideation necessitating immediate treatment was also exclusionary. All subjects accepted into the study were strongly right-handed with a score of 35 or greater on the 39-point scale of handedness (Chapman and Chapman, 1987). Figure 1 provides a flow chart of the participant screening and recruitment process; for more information see (Stewart et al., 2010). Seven participants were excluded from the final statistical analysis because of the presence of unrecoverable ectopic beats in their EKG data. Another twelve participants were excluded because of an excess of artifact-dominated segments in their EEG data. The final sample for statistical analyses consisted of 287 participants. The lifetime MDD group (N=134, 38 male, 77.55 % Caucasian, 21.94 % Hispanic) comprised participants who met DSM-IV criteria for past or current major depressive episode. Participants without any Axis I disorder were considered the healthy control group (N=153, 51 male, 71.62 % Caucasian, 25.47 % Hispanic). For more details about subjection recruitment procedures and the demographic information, see (Stewart et al., 2010).

**Figure 1:**
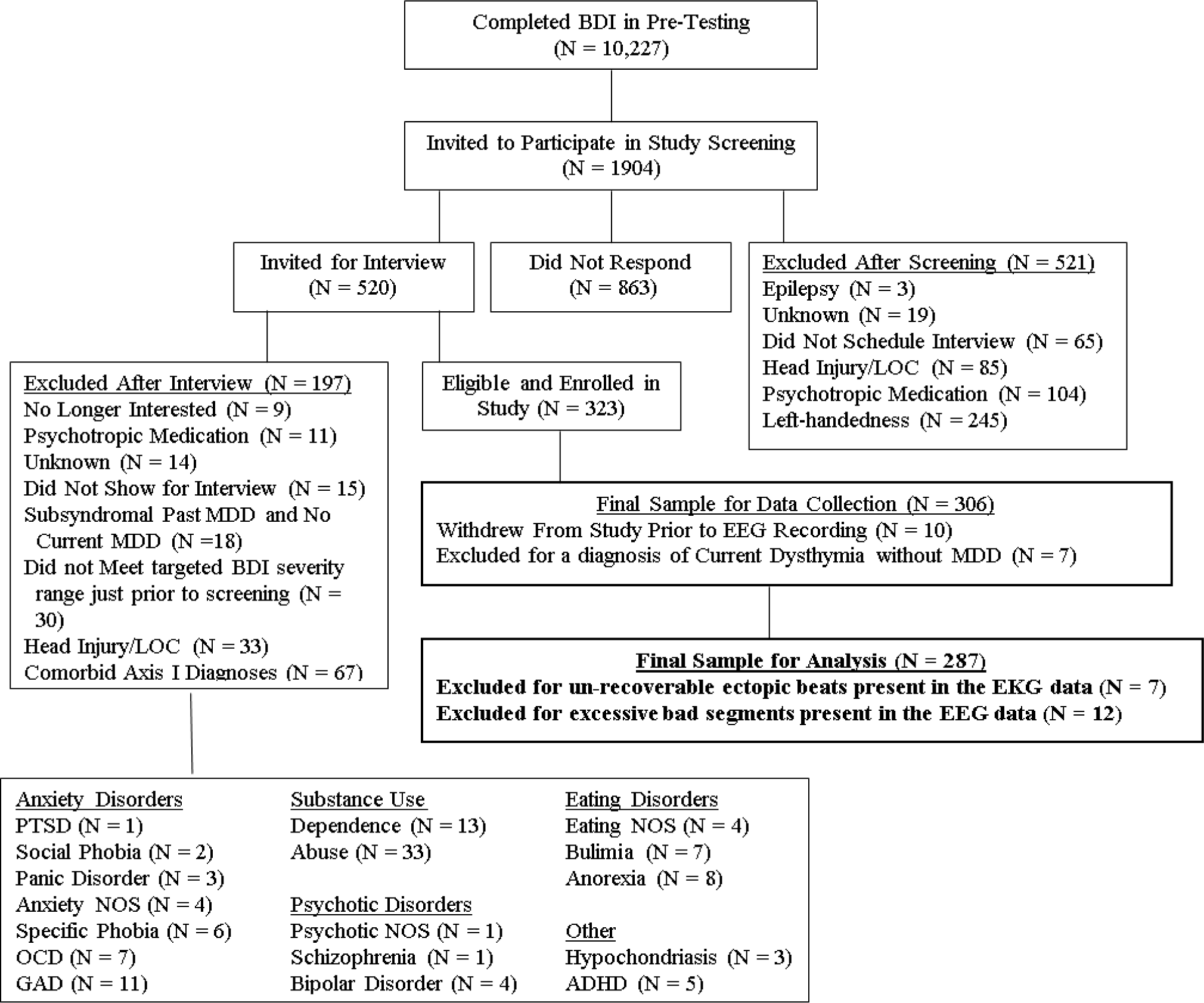
Flowchart of participant screening and enrollment procedure. BDI=Beck Depression Inventory; LOC=loss of consciousness; MDD=major depressive disorder; EEG=electroencephalographic; PTSD=post-traumatic stress disorder; NOS=not otherwise specified; OCD=obsessive compulsive disorder; GAD=generalized anxiety disorder; ADHD=attention deficit hyperactivity disorder. Flowchart modified from (Stewart et al., 2010)

### 2.2 | Simultaneous EEG & EKG Data Collection

All participants provided informed consent prior to EEG/EKG data collection. Data collection was conducted on four separate days with at least 24 hours between visits and with all four visits completed within a period of two weeks. There were two simultaneous resting EEG/EKG sessions at each visit, separated by approximately 20 minutes. Each session consisted of eight 1-minute resting data collection blocks with either eyes closed (C) or eyes open (O), in one of two counterbalanced orders (OCCOCOOC or COOCOCCO). The recording system was a 64-channel NeuroScan Synamps2 amplifier and acquisition system (Neuroscan, 2008), with a one G ohm input impedance and a 16-bit analog-to-digital converter sampling at 1000 Hz. Ag-AgCl electrodes were used, with impedances kept lower than 10 Kohms, and placed according to the International 10-20 system (Homan et al., 1987). Two electrooculogram (EOG) channels (vertical EOG and horizontal EOG) were added for ocular artifact rejection. Both EEG and EKG data were filtered with a 200 Hz low-pass filter, amplified 2816 times and then digitized at 1000 Hz. The ground electrode was located anterior to Fz, while the online reference electrode was immediately posterior to Cz. More information about the data collection procedure can be found in (Stewart et al., 2010).

### 2.3 | EKG Data Preprocessing and Reduction

Eight blocks of one-minute EKG data (Panel A in Figure 2) were recorded concurrently with the EEG data in a Lead I configuration with two Ag/AgCl sensors attached to the collarbones. The EKG signal was amplified 2816 times and sampled at 1000 Hz. R-spikes were identified in QRSTool (Allen et al., 2007), an EKG beat detection software program that allows for modification of beat assignments while simultaneously showing the resulting inter-beat interval (IBI) time series (Panel B in Figure 2). To avoid missed or extraneous beats, all EKG files were subsequently scored by human raters. The IBI series was then converted into a time-series and filtered using an optimal finite impulse response filter to extract the activity in the respiratory frequency band (.12-.4 Hz; Panel C in Figure 2). The Hilbert transform (Hahn, 1996; Huang et al., 2009) was then applied to the filtered IBI time series. The amplitude of the transformed signal (Panel D in Figure 2) represents the instantaneous variability of the filtered IBI time series. A more conventional moving window approach (window size=10 s, offset=1 s) was also applied to the filtered IBI to derive moving window RSA (Panel E in Figure 2). Figure 2 shows that Hilbert RSA (Panel D) tracks the variability in the filtered IBI (Panel C) better than moving window RSA (Panel E). Thus, the Hilbert RSA signal was inserted into the EEG data structure and aligned with the timing of the EEG channels and later used for splitting epochs into above vs below median RSA categories. There’s no need to compare Hilbert transformed RSA with moving-widow RSA.

**Figure 2:**
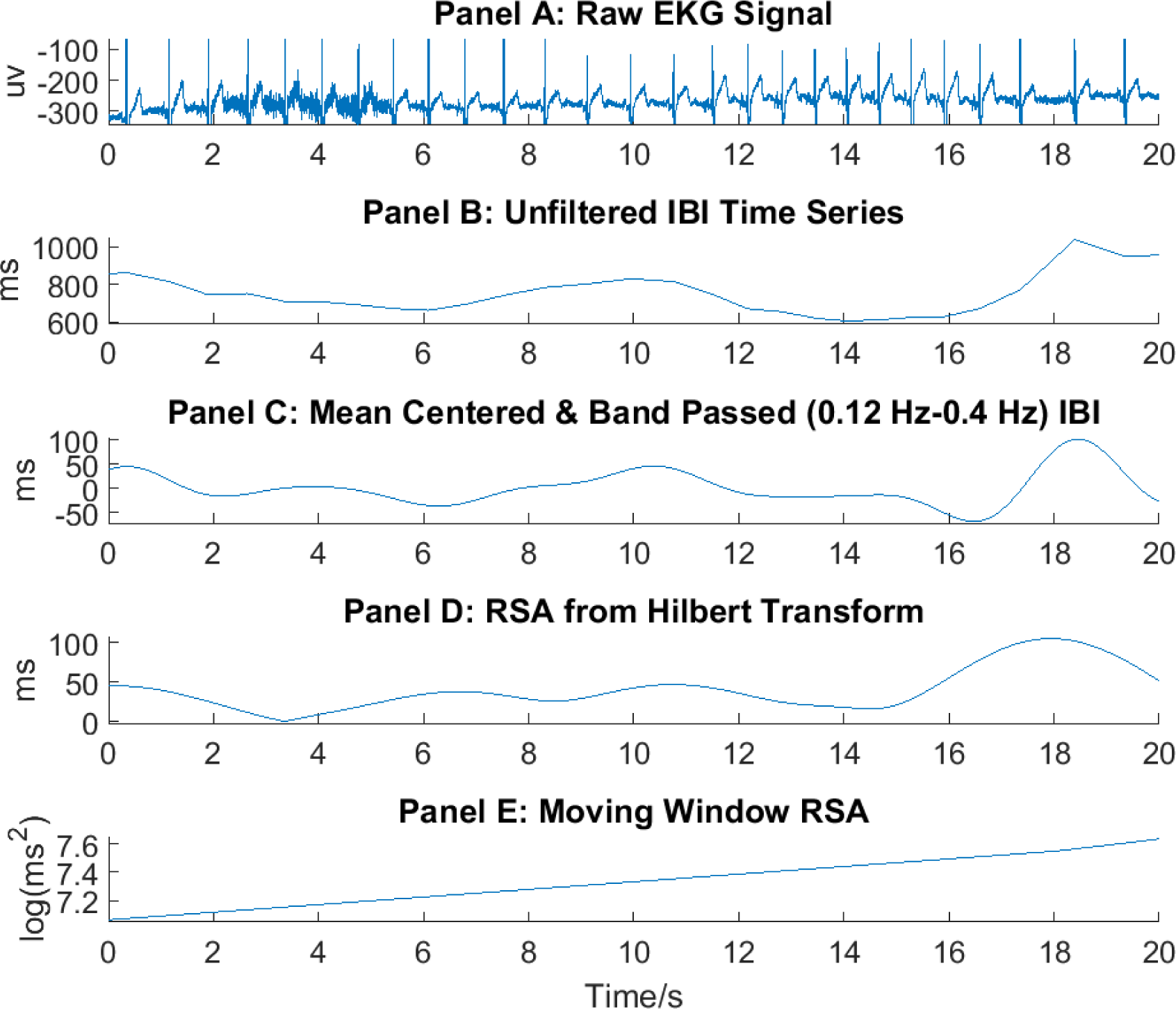
Steps involved in the preprocessing & reduction of EKG data. (20 seconds of data depicted). All the x axes have the same label and units, i.e., Time/s. IBI=Inter-beat Interval; RSA=Respiratory Sinus Arrhythmia, which is thought to index vagally-mediated heart rate variability; Moving Window= Moving Window approach (window size=10s, offset=1s)

### 2.4 | EEG Data Preprocessing and Reduction

All EEG data were visually inspected by trained human raters to remove movement and muscle artifacts before preprocessing. EEG data preprocessing and reduction were implemented using custom scripts written in Matlab (The Mathworks, Inc., Natick, MA) with various functions adapted from the EEGLab (Delorme and Makeig, 2004). The preprocessing included the following steps in the order of their implementations. Step 1: Remove any bad data segment identified from visual inspection prior to data preprocessing. Step 2: Divide each one-minute data block into 2.048-second epochs with 75% overlap between consecutive epochs (to compensate minimal weights applied to the end of each epoch when Hamming window is applied). Step 3: Scan each epoch for ocular artifacts and reject any epoch with greater or less than +/- 75 microvolt ocular activity. Step 4: Reject epochs with data points that were 6 standard deviations above or below the mean of the whole epoch. Step 5: Take the current source density transform (Kayser and Tenke, 2015) of each classified EEG epoch, using the laplacian_perrinX function from (Cohen, 2014) with default settings. Step 6: Apply a Hamming window function (to reduce edge artifacts) to each epoch of CSD transformed data and then take the Fast Fourier Transform of each epoch. Step 7: Separately for eyes open resting epochs and eyes-closed resting epochs, find the median value (M1) within each epoch of RSA derived by the Hilbert method; find the median value (M2) across epochs of these median values (M1s). Step 8: Also separately for eyes open and closed, take the mean of power spectra for those epochs above the median (M2) and for those below the median (M2). Step 8: Extract the total alpha power (8-13 Hz) from the averaged power spectra for each electrode and each condition (eyes open/closed by above/below median). Step 9: Take the log transform of the averaged alpha power at each electrode site and calculate the right-left difference scores at four homologous pairs of electrodes (i.e.: F2-F1, F4-F3, F6-F5, F8-F7) to get the frontal alpha asymmetry scores. Because alpha has an inhibitory effect on cortical network activity (Allen, Coan, Nazarian, 2004), higher scores reflecting relatively greater right frontal alpha are interpreted as indexing relatively greater left frontal activity.

### 2.5 | Statistical Analyses

The final dataset for statistical analysis included asymmetry scores at 4 frontal electrode pairs (F2-F1, F4-F3, F6-F5, and F8-F7), on 4 days, for eyes open versus closed, and for epochs where Hilbert RSA was above and below the median within the eyes open or eyes closed epochs. Between subject variables included lifetime MDD status and sex. Within-subject variables were frontal electrode pair, open/closed resting condition, day, and Hilbert RSA condition (above/below median). Using the “lmer” function from the “lme4” package (Bates et al., 2014) in R (Team, 2014), a linear mixed model was fitted to the data via the restricted maximum likelihood estimation method, with the resting front alpha asymmetry score (rLFA) as the dependent variable; Hilbert RSA, lifetime MDD status (Lifetime_MDD), sex, electrode pair (Elect_pair), eyes open or closed resting condition (Rest), the interaction between Hilbert RSA and lifetime MDD, the interaction between Hilbert RSA and electrode pair, the interaction between Hilbert RSA and sex, and the interaction between Hilbert RSA and resting condition as the fixed effects. A correlated random effect of Hilbert RSA by Day was included in the model to account for the fact that the effect of RSA on rLFA could vary randomly across the four days when data were collected.

## 3 | RESULTS

The overall model predicting relative left frontal activity has a total explanatory power (conditional *R*^2^) of 6%, in which the fixed effects explain 5.7% of the variance (marginal *R*^2^). Relating to the first hypothesis, there was a main effect of Hilbert RSA on rLFA (F(1, 18)=41.9041, p<0.0001) such that when Hilbert RSA was lower, rLFA score was also lower (See Table 2 & Figure 3). Regarding the prediction from our second hypothesis, this association of Hilbert RSA and rLFA was not found to be moderated by Lifetime MDD status (F(1,16030)=1.0720, p=0.3005 (See Table 2 & Figure 4). However, an interaction effect of RSA and electrode pair on rLFA was found (F(3, 16030)=46.8273, p<0.0001; See Table 2 & Figure 5). Specifically, the main effect of RSA on rLFA was in the predicted direction only for F3_F4 and F8_F7, but not for F2_F1 or F6_F5, with the latter two electrode pairs showing a small effect in the direction contrary to prediction from Hypothesis I (see Figure 5). Regarding the other control variables that were included in the model, the main effect of RSA on rLFA was not moderated by sex (F(1,16030)=2.2252, p=0.1358; see Table 2 & Figure 6) or eyes open/closed conditions (F(1, 16030)=0.7188; p=0.3966; see Table 2). In addition to the two main hypotheses that we tested, we also replicated the main effects of lifetime MDD status, sex, resting condition and electrode pair on alpha asymmetry that were previously reported in (Stewart et al., 2010). Finally, regarding the correlated random effect of Hilbert RSA by Day, we did not find it significant (LRT=0.4197, df=2, p=0.8107; see Table 3). That is, the effect of RSA on rLFA did not vary across day in a statistically significant fashion (see Figure 8).

**Figure 3:**
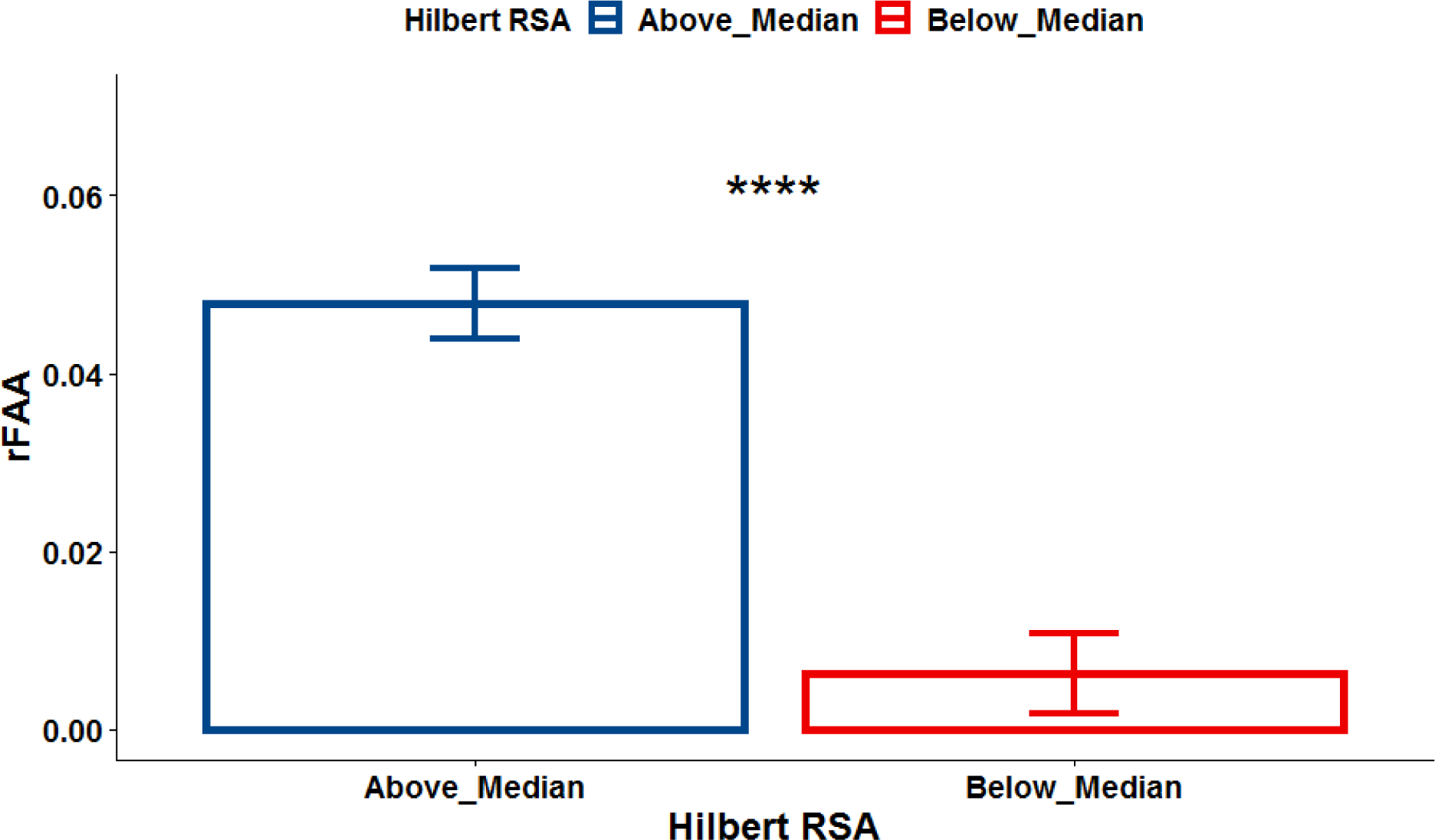
Within-Subject Association of rLFA and RSA. Note: rLFA=relative left frontal activity; RSA=Respiratory Sinus Arrhythmia; Above_Median=Epochs with Above Median Hilbert RSA; Below_Median=Epochs with below median; Hilbert RSA= Hilbert Transformed RSA. The Wilcox test is used for the post-hoc comparisons. Error bars represent standard errors of the estimates. ****=p<0.0001, ***=p<0.001, **=p<0.01, *=p<0.05.

**Figure 4:**
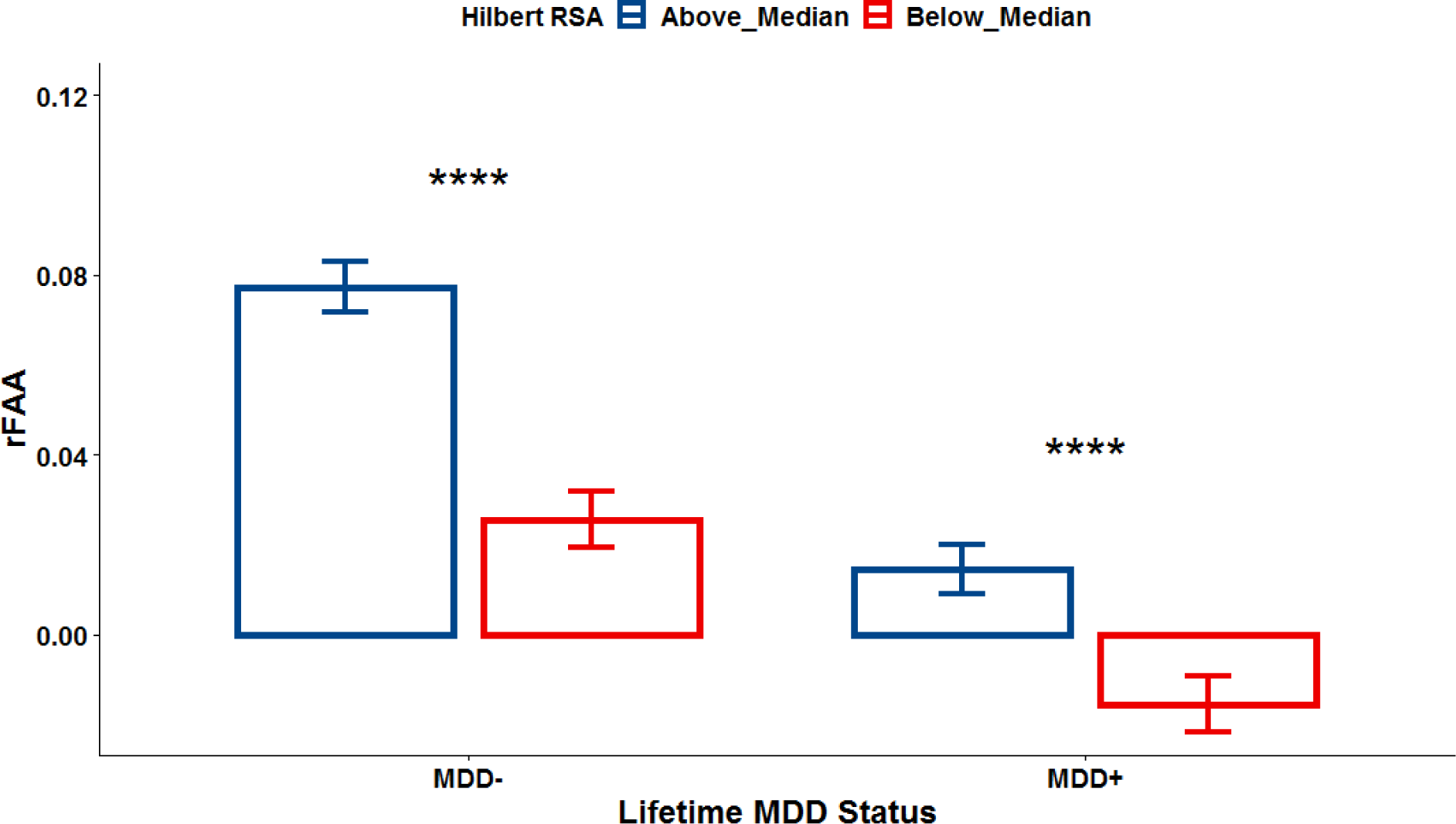
Within-Subject Association of rLFA and RSA Not Moderated by Lifetime MDD. Note: rLFA=relative left frontal activity; RSA=Respiratory Sinus Arrhythmia; Above_Median=Epochs with Above Median Hilbert RSA; Below_Median=Epochs with below median; Hilbert RSA =Hilbert Transformed RSA; MDD-=Subjects without a history of MDD; MDD+=Subjects current diagnosed with MDD or with a history of MDD. The Wilcox test is used for the post-hoc comparisons. Error bars represent standard errors of the estimates. ****=p<0.0001, ***=p<0.001, **=p<0.01, *=p<0.05.

**Figure 5:**
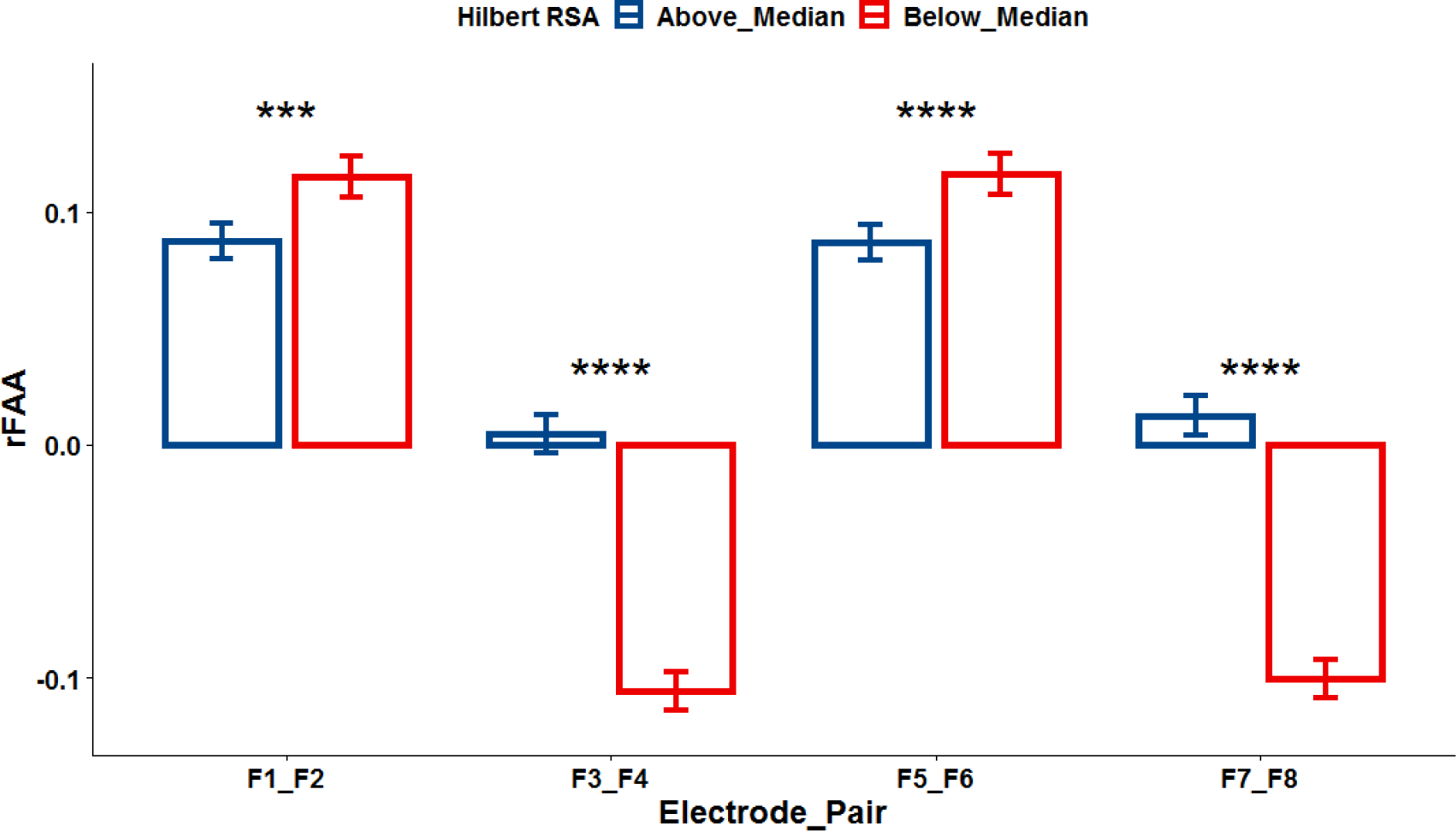
Within-Subject Association of rLFA and RSA Moderated by Electrode Pair. Note: rLFA=relative left frontal activity; RSA=Respiratory Sinus Arrhythmia; Above_Median=Epochs with Above Median Hilbert RSA; Below_Median=Epochs with below median; Hilbert RSA= Hilbert Transformed RSA. The Wilcox test is used for the post-hoc comparisons. Error bars represent standard errors of the estimates. ****=p<0.0001, ***=p<0.001, **=p<0.01, *=p<0.05.

**Figure 6:**
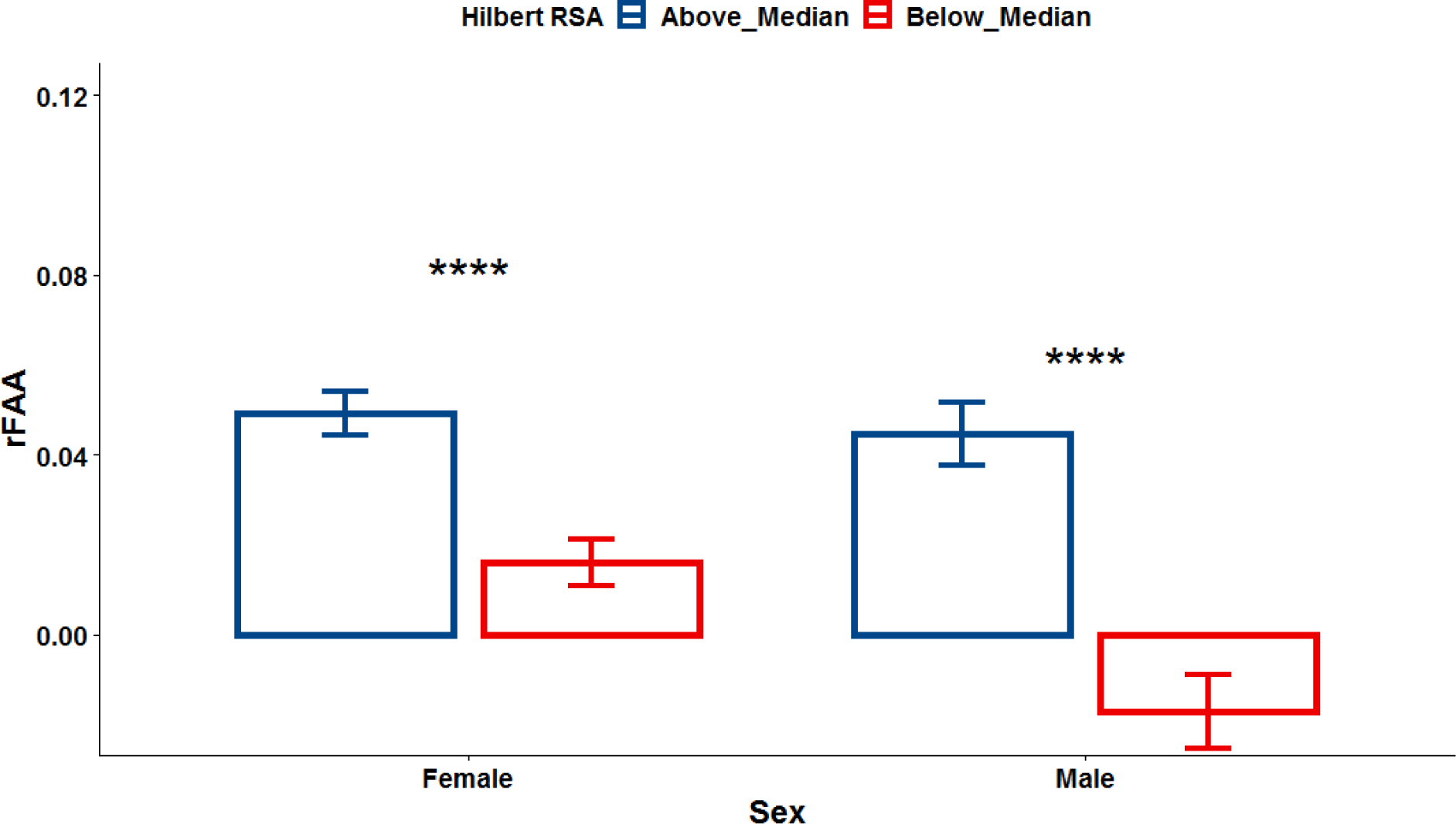
Within-Subject Association of rLFA and RSA Not Moderated by Sex. Note: rLFA=relative left frontal activity;, RSA=Respiratory Sinus Arrhythmia;, Above_Median=Epochs with Above Median Hilbert RSA;, Below_Median=Epochs with below median;, Hilbert RSA= Hilbert Transformed RSA. The Wilcox test is used for the post-hoc comparisons. Error bars represent standard errors of the estimates. ****=p<0.0001, ***=p<0.001, **=p<0.01, *=p<0.05.

**Figure 7:**
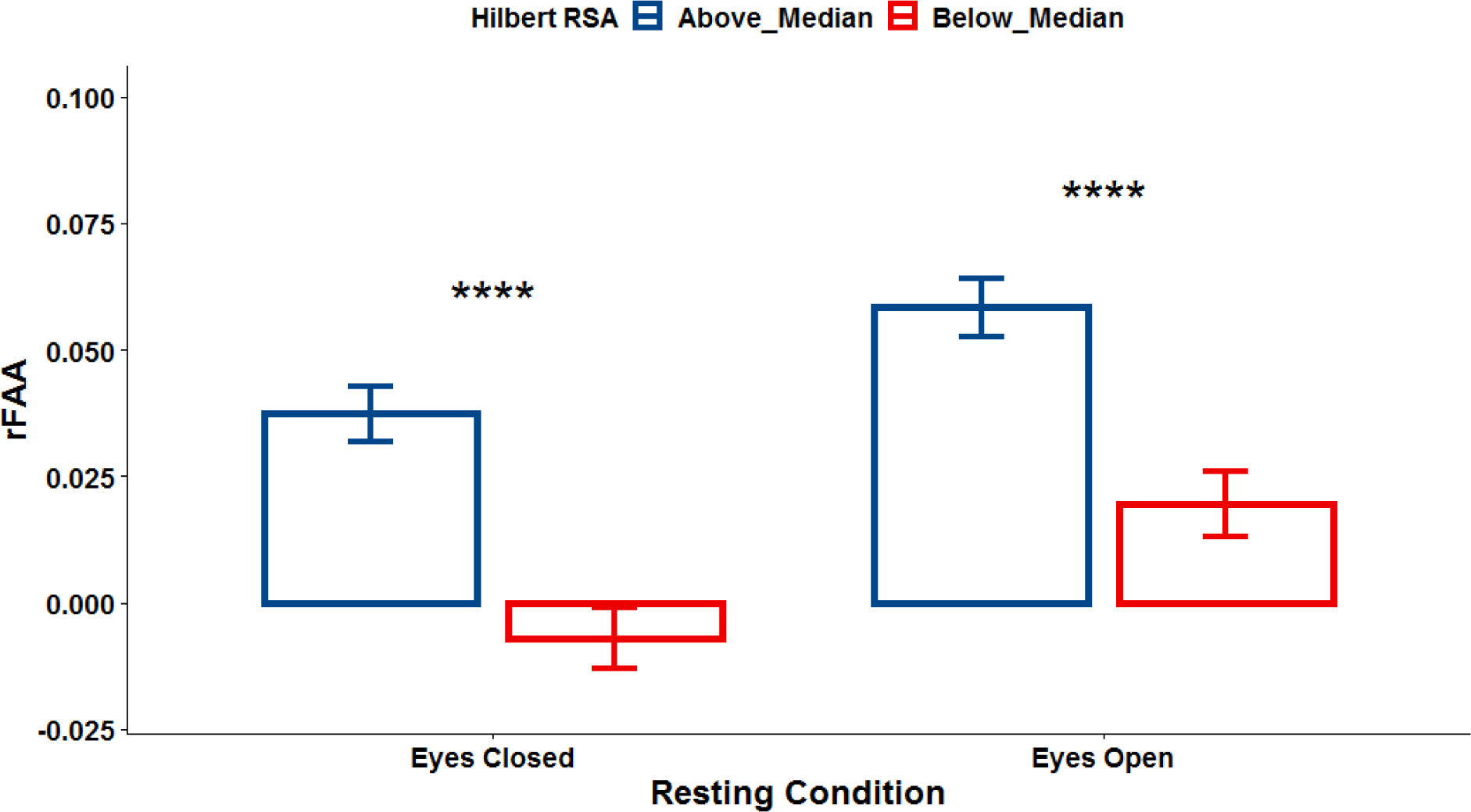
Within-Subject Association of rLFA and RSA Not Moderated by Eyes Open/Closed. Note: rLFA=relative left frontal activity;, RSA=Respiratory Sinus Arrhythmia;, Above_Median=Epochs with Above Median Hilbert RSA;, Below_Median=Epochs with below median;, Hilbert RSA= Hilbert Transformed RSA. The Wilcox test is used for the post-hoc comparisons. Error bars represent standard errors of the estimates. ****=p<0.0001, ***=p<0.001, **=p<0.01, *=p<0.05.

**Figure 8:**
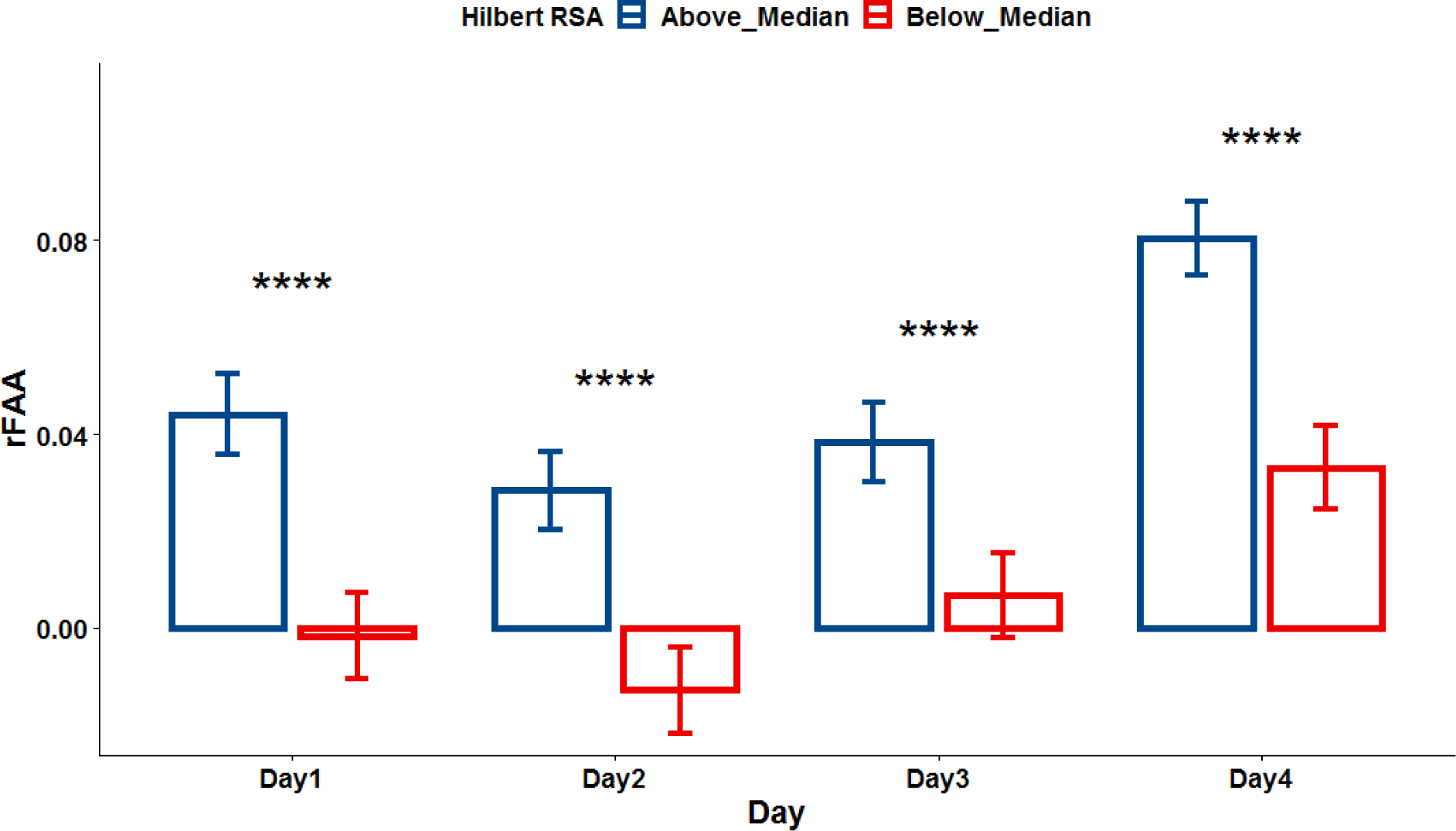
Within-Subject Association of rLFA and RSA Not Moderated by Eyes Open/Closed. Note: rLFA=relative left frontal activity;, RSA=Respiratory Sinus Arrhythmia;, Above_Median=Epochs with Above Median Hilbert RSA;, Below_Median=Epochs with below median;, Hilbert RSA= Hilbert Transformed RSA. The Wilcox test is used for the post-hoc comparisons. Error bars represent standard errors of the estimates. ****=p<0.0001, ***=p<0.001, **=p<0.01, *=p<0.05.

**Table 1.**
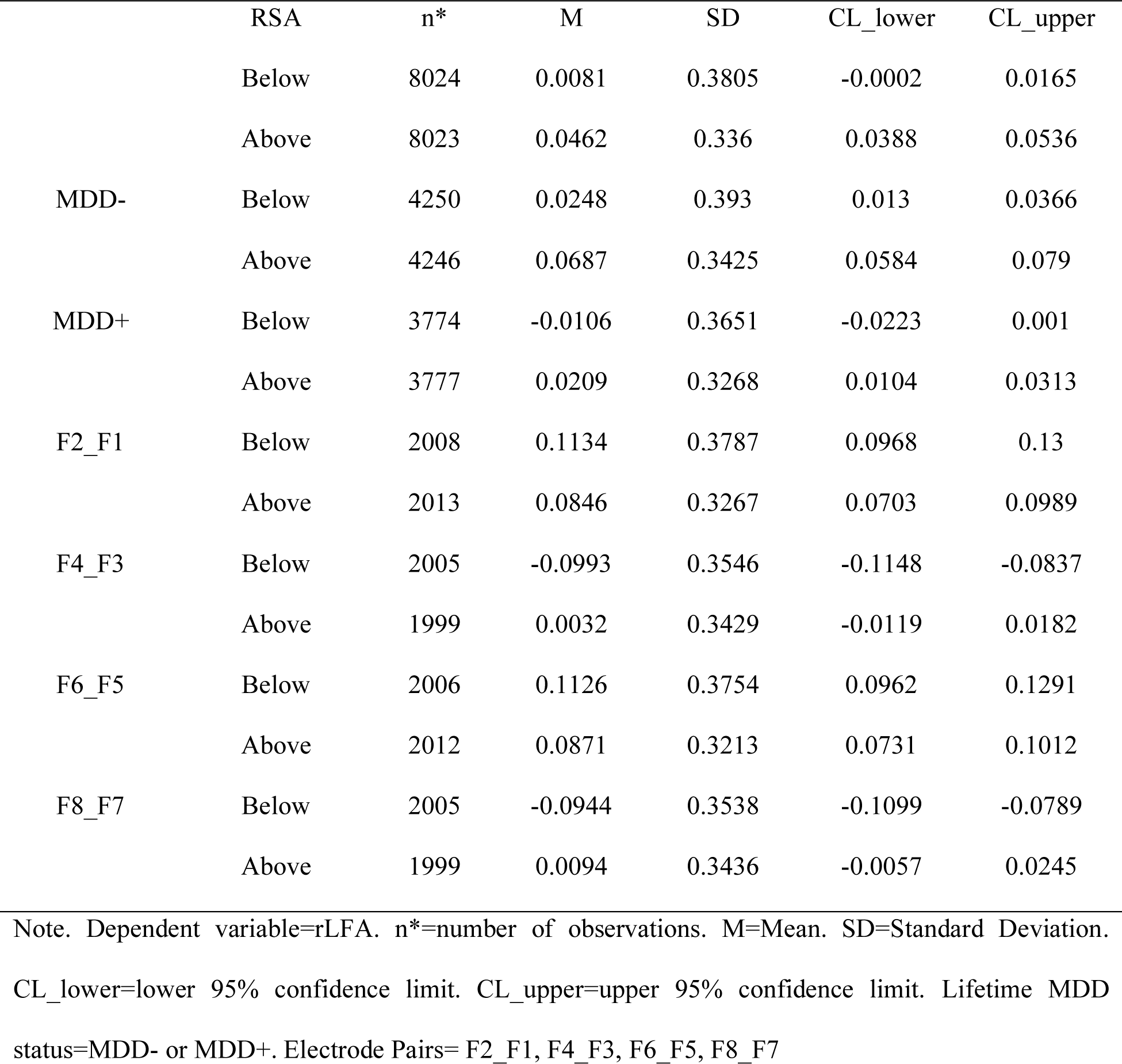
*Descriptive Statistics*

**Table 2.**
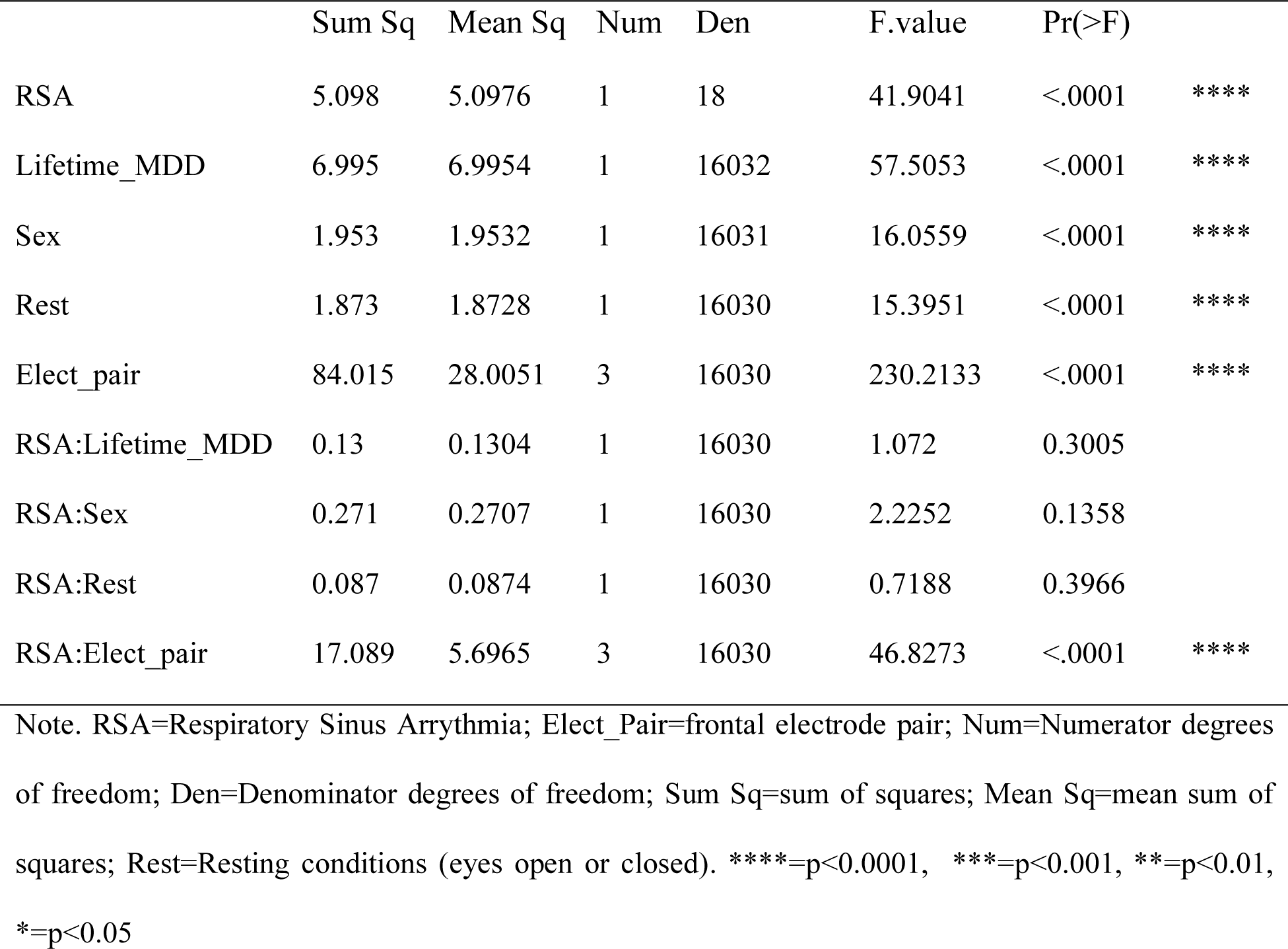
*Analysis of Variance Table of type III with Satterthwaite approximation for degrees of freedom*

**Table 3.** *ANOVA-like table for random-effects: Single term deletions*

|  | npars | logLik | AIC | LRT | Df | Pr(>Chisq) |
| --- | --- | --- | --- | --- | --- | --- |
| Random effect | 18 | -5919.1 | 11874 |  |  |  |
| (RSA\|Day) | 16 | -5919.3 | 11871 | 0.41967 | 2 | 0.8107 |

*Note.* npars=number of model parameters, logLik=the log-likelihood for the model or REML log-likelihood, AIC=Akaike Information Criterion, LRT=the likelihood ratio test statistics; twice the difference in log-likelihood, which is asymptotically chi-square distributed, Df=degrees of freedom for the likelihood ratio test, Pr(>Chisq)=the p-value.

## 4 | DISCUSSION

This is the first study investigating the within-person relationship between rLFA resting alpha asymmetry and RSA in participants with a history of depression. An association between rlFA and RSA was found, such that during periods when individuals had relatively lower RSA, they had relatively less left frontal activity (indicated by lower alpha asymmetry score). This association is interesting because both lower RSA (Koch et al., 2019; Thayer et al., 2012) and relatively less left frontal activity (lower rLFA scores) (Sutton and Davidson, 1997) have been observed in participants with a history of depression and linked to withdrawal motivation and poorer emotional regulation capacity. The finding is also consistent with the prediction from the homeostatic neuroanatomical model of forebrain emotional asymmetry (Craig, 2005) . Regarding our second hypothesis, however, we did not find a significant interaction between Hilbert RSA and lifetime depression status on relative left frontal activity. This association of RSA and rLFA was not moderated by sex or resting condition (eyes open or closed). Taken together, it seems that this association of RSA and alpha asymmetry is quite robust. However, the within-person association seems to be moderated by the specific frontal electrode pairs. One possible explanation might be that the frontal cortex is large and functionally heterogeneous. The electrode pairs F2_F1, F4_3, F6_F5, and F8_F7 indeed cover both the lateral and medial portions of the prefrontal cortex, and the CSD transformation used here will accentuate the local contributions at each recording site. Further research is needed to determine the spatial extent of this effect, e.g., examining the relationship between source-localized EEG and RSA to see whether there are specific cortical regions that are sensitive to changes in vagal control of the heart.

Another limitation of the present study is that the directionality of the association between resting frontal EEG alpha asymmetry and vagally-mediated RSA cannot be obtained from the present findings. Data were aligned in real time, although the impact of central nervous system (CNS) efferent activity on cardiac timing will not be manifest until the subsequent cardiac cycle. Given the 2.048 sec epoch length, the impact of CNS activity within the epoch may often be reflected in the timing of the IBIs still within that epoch, and the Hilbert-transformed band-limited IBI series should provide a near instantaneous estimate of that effect. Moreover, given the overlapping epochs used for extracting spectral power with hamming-windowed EEG data, adjacent epochs are likely to be highly similar, and captured by the median-split approach used here. Future work may profit from examining lagged covariance of these signals, but this was not possible with the present data given that EEG artifact removal resulted in gaps (excluded epochs) in the continuous signals. Nonetheless, the present findings suggest that there is within subject association of a central nervous system marker (EEG alpha asymmetry) and an autonomic nervous system marker (RSA) of depression, consistent with findings using concurrent EKG and fMRI data (Schafer et al., 2015).

If in future studies, source-localized EEG activity from the left frontal cortex leads changes in RSA, it would lend support for the homeostatic neuroanatomical model of forebrain emotional asymmetry (Craig, 2005). On the other hand, if increases in RSA lead increases in left frontal activity (as measured by source-localized EEG), this would suggest that autonomic state may have an important influence on cortical activity and information processing. If this were indeed the case, interventions designed to enhance vagal control (e.g. paced breathing) could produce changes in brain activity and mood. However, we speculate that the influences between left frontal activity and RSA are most likely bi-directional; i.e., left frontal activity can both lead or lag changes in RSA. This would be in accordance with both the homeostatic neuroanatomical model of forebrain emotional asymmetry (Craig, 2005) and the model of neurovisceral integration (Smith et al., 2017; Thayer and Lane, 2000, 2009; Thayer et al., 2012). According to the neurovisceral integration model, the frontal cortex influences and is influenced by cardiac activity via direct and indirect pathways linking the frontal cortex to the Central Autonomic Network (CAN)–a network of brain regions that produces sympathoexcitatory and parasympathoinhibitory effects on the heart (Thayer and Lane, 2009). Future studies might tease apart these directional effects by manipulating vagal control (e.g., vagal stimulation, paced breathing) or manipulating frontal EEG asymmetry (cf. Allen et al., 2001; Harmon-Jones, 2006) to assess the impact frontal EEG asymmetry and RSA respectively.

## Author Notes

The authors wish to thank Jennifer Stewart, Jim Coan, Dave Towers, Jamie Velo, and Johnny Vanuk for assistance with data acquisition and processing. This study was funded in part by grants from the National Institute of Mental Health (grant R01–MH066902 and R21-MH101398), and from the National Alliance for Research on Schizophrenia and Depression (NARSAD). Address correspondence to John JB Allen, Department of Psychology, University of Arizona, P.O.Box 210068, Tucson, AZ, 85721-0068. Email

